# Tendon-specific PIEZO1 activation enhances neural drive to muscle during ballistic movement

**DOI:** 10.64898/2026.11.07.765061

**Authors:** Kohei Nakamura, Akito Kosugi, Ryo Nakamichi, Ryota Kurimoto, Takahide Matsushima, Tomoki Chiba, Masatoshi Ko, Takahiro Morita, Toshitaka Yoshii, Kazuhiko Seki, Hiroshi Asahara

## Abstract

Tendons are generally regarded as passive structures that transmit muscle force to bone, yet they contain the Golgi tendon organ (GTO), whose Ib afferents act on motor neurons either inhibitory or excitatory depending on movement context. Whether a change in tendon mechanical properties alters the neural drive to muscle during movement remains unknown. We therefore recorded electromyography (EMG) and ankle kinematics during jumping in male mice carrying a tendon-specific gain-of-function (GOF) mutation in the mechanosensitive channel PIEZO1, which renders the tendon more compliant. In wild-type (WT) mice, gastrocnemius EMG amplitude increased with jump distance but not tibialis anterior activity, identifying the gastrocnemius as the muscle that scales with jump demand. At their maximum jump distance, which was longer in GOF than in WT mice, GOF mice showed a larger gastrocnemius EMG amplitude with unchanged burst duration. No genotype difference was detected at a submaximal distance, indicating that this enhancement depends on load. Kinematics showed the same specificity: GOF mice reached a deeper peak dorsiflexion before takeoff only at maximum jump distance, and the EMG difference emerged only after maximal dorsiflexion, not at burst onset. Muscle mass and myogenic gene expression were unchanged, whereas tendon marker genes were elevated, indicating that the enhancement was not of muscular origin. These findings indicate that a change in tendon mechanical properties alters the neural drive to muscle in a load-dependent manner, most plausibly through positive force feedback from the GTO, and identify the tendon as a sensory organ whose mechanical properties shape motor output.

**Significance Statement:** Tendons are usually treated as passive cables that transmit muscle force to bone, and studies of motor control have focused on muscle and neural circuits. Here we show that a genetic modification confined to tendon changes how strongly muscle is recruited during jumping in mice. Tendon-specific *Piezo1* gain-of-function mice generated larger gastrocnemius activity than controls, but only during maximal jumps, and this difference appeared after the ankle had reached its maximal dorsiflexion. Muscle mass and myogenic gene expression were unchanged. These results suggest that the tendon is not merely a passive force transmitter but a sensory organ whose mechanical properties shape the neural drive to muscle, offering a new framework for understanding motor performance and its decline with tendon aging.

## Introduction

Tendons connect muscle to bone and transmit muscle force during movement (Benjamin et al., 2008). Although tendon has long been regarded as a stable tissue with minimal cellular and matrix turnover (Heinemeier et al., 2013), its collagen-based mechanical properties are not fixed after birth: they change with development, training (Langberg et al., 2001; Bohm et al., 2015), and aging (Heinemeier et al., 2013; Stowe et al., 2024). These changes have a large effect on motor output: age-related tendon stiffening is associated with a decline in motor function (Nakamichi and Asahara, 2024), whereas tendons with greater compliance improve jumping performance (Nakamichi et al., 2022). This effect has been explained primarily in mechanical terms (Hill, 1938): because tendon is arranged in series with muscle, a more compliant tendon stores and releases more elastic energy (Roberts and Azizi, 2011; Nakamichi et al., 2022) and allows muscle fascicles to shorten at a more favorable length and velocity (Kurokawa et al., 2001), thereby directly increasing muscle power output.

Tendon could also influence motor output by a second, indirect route through the central nervous system. Tendons contain the Golgi tendon organ (GTO), a force receptor at the muscle–tendon junction whose Ib afferents signal tendon tension to the spinal cord (Stuart et al., 1972; Jami, 1992). Because the GTO lies in series within the muscle–tendon unit, the strain it experiences at a given muscle force depends on the compliance of the surrounding collagen (Jami, 1992); a change in tendon mechanical properties should therefore alter the Ib signal the spinal cord receives for the same muscle force. Ib input from the GTO has conventionally been understood as autogenic inhibition, in which Ib afferents inhibit motor neurons of the homonymous muscle and thereby protect it from overload (Pearson and Collins, 1993; Nielsen and Jessell, 2021). However, during locomotion this pathway reverses to excitation and provides positive force feedback that reinforces extensor activity (Guertin et al., 1995; Donelan et al., 2009), with a gain that depends on both tendon force and the length of the muscle–tendon unit (Donelan and Pearson, 2004). We recently showed that this excitatory Ib feedback loop also contributes to the generation of muscle activity during voluntary movement in primates (Kim et al., 2026).

These observations predict that a change in tendon mechanical properties alters the Ib input to this circuit and thereby the neural drive to muscle. Whether this occurs, and how it shapes motor output, has not been examined, largely because the interventions used to alter tendon properties, such as training, unloading by hindlimb suspension, and aging, change the muscle and the tendon at the same time (Heinemeier et al., 2013; Bohm et al., 2015), so that neural changes could not be attributed to the tendon alone.

We recently developed a mouse in which a gain-of-function (GOF) mutation of the mechanosensitive channel PIEZO1 is expressed only in tendon under the *Scx*-Cre driver (Nakamichi et al., 2022). In these mice the tendon is remodeled and more compliant, and jumping performance is improved, whereas muscle mass, fiber-type composition, and GTO morphology are unchanged, and the same mutation introduced into muscle does not improve performance (Nakamichi et al., 2022). Neural function in this line, however, has not been assessed. Furthermore, in that report the increase in kinetic energy during jumping far exceeded the increase in elastic energy stored in the tendon, indicating a contribution from a mechanism other than tendon energy storage (Nakamichi et al., 2022), such as a change in neuromuscular activation.

Here we used these mice to examine whether a tendon-restricted change in mechanical properties alters the neural drive to muscle during movement. We recorded hindlimb EMG and ankle kinematics during jumping and compared the two genotypes. Because Ib input scales with tendon force (Donelan and Pearson, 2004), we examined jumps of different effort levels, asking whether the influence of the altered tendon on neural drive depends on the load imposed on the muscle–tendon unit.

## Materials and methods

### MICE

Using the Cre/loxP system, *Piezo1*cx (gain-of-function allele) mice were crossed with *Scx*-Cre mice, and *Scx*-Cre:*Piezo1*cx/+ mice were used as tendon-specific *Piezo1* GOF mice (GOF mice) (Fig. S1). Littermate *Scx*-Cre-negative *Piezo1*cx/+ male mice were used as controls (WT). All mice used were 12-week-old males. All animal procedures were performed in accordance with the Policies on the Use of Animals and Humans in Research. All mice were bred and maintained at the No. 1 Animal Housing Facility, Animal Research Center, Institute of Science Tokyo, and were handled in accordance with an animal experimentation protocol (Protocol No. A2026-047C2) approved by the Institutional Animal Care and Use Committee of Institute of Science Tokyo.

### EXPERIMENTAL DESIGN

The purpose of this study was to determine whether neural activity during jumping is altered in tendon-specific *Piezo*1 GOF mice, and thus whether the tendon can shape neuromuscular output. The experimental design is outlined below (Fig. 1A). On experimental Day 0, mice were moved from the housing room to the experimental room. On Day 1, a jump test and EMG electrode implantation surgery were performed. On Day 3, the wound and electrode connector were checked. On Day 4, EMG during jumping was measured, followed by CMAP measurement. On Day 8, tissue sampling was performed. First, EMG was measured in a stepwise manner at 15 cm, 25 cm, and 35 cm in a single WT mouse to confirm the correlation between hindlimb muscle EMG and jump distance within the same individual. Next, EMG waveforms at maximum jump distance in WT and GOF mice were normalized using CMAP data and compared between groups. Seventeen WT mice and 15 GOF mice were used for electrode implantation and EMG analysis.

**Figure 1.**
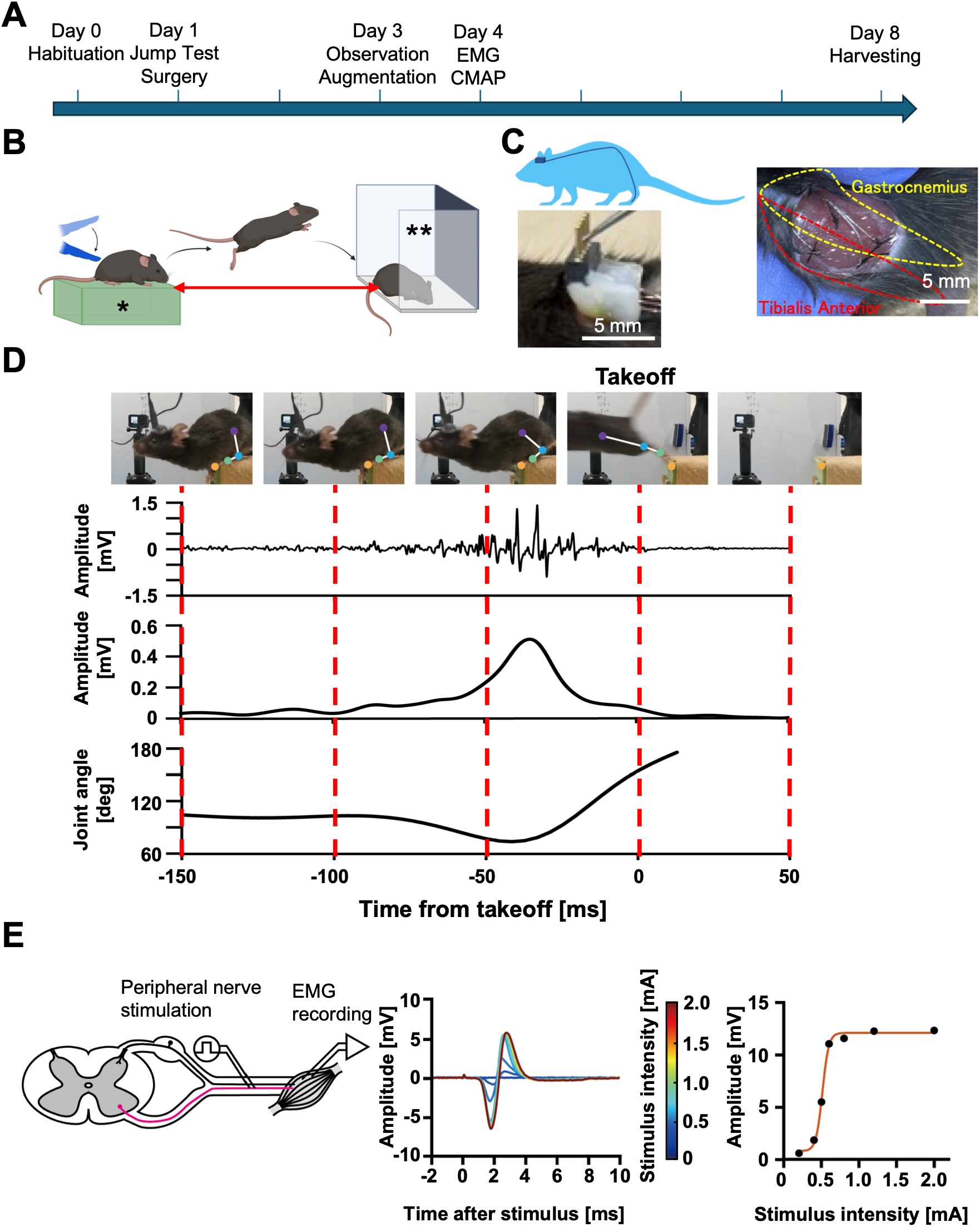
Experimental Method. (A) Experimental workflow. (B) Schematic of the jump test. The mouse is placed on the platform and prompted to jump across to the cage by a tactile stimulus applied to the base of its tail. A single asterisk (*) denotes the platform, and a double asterisk (**) denotes the cage. The red double-headed arrow indicates the jump distance. (C) The top left panel shows the placement of the head connector and the subcutaneous route of the electrode wires. The bottom left panel shows the EMG connector fixed to the skull. The right panel shows the gastrocnemius and tibialis anterior, into which the electrodes were inserted. (D) Representative single jump from one WT mouse. Signals were aligned to takeoff (t = 0 ms), defined as the first frame in which both hindlimbs had left the platform. Top row, video frames at the times marked by the dashed lines. Colored dots denote tracked landmarks: purple, knee; blue, ankle; green, foot; orange, beam 1 (the distal edge of the platform). Second row, raw gastrocnemius EMG shown from 150 ms before to 50 ms after takeoff. Third row, the EMG envelope of the same signal. Fourth row, ankle joint angle shown from 150 ms before to 12.5 ms after takeoff. (E) Normalization of the EMG amplitude to the maximum compound muscle action potential (CMAP). The left panel shows the schema of the in vivo electrophysiological experiment. Electrical stimulation was applied to the sciatic nerve under anesthesia, and the evoked CMAPs were recorded. The middle panel shows examples of CMAPs at different stimulus intensities. The right panel shows the change in CMAP amplitude as a function of stimulus intensity, fitted with a four-parameter logistic function.

### JUMP TEST

The experimental setup consisted of a wooden jump platform (3 cm wide, 20 cm long, Fig. 1B) on a table 80 cm high, and a cage (15 cm high, 25 cm deep, 50 cm wide, Fig. 1B). Mice were placed in the cage and allowed to habituate for 5 minutes before being moved onto the jump platform. At a platform-to-cage distance (jump distance, Fig. 1B) of 5 cm, a tactile stimulus was applied to the base of the mouse’s tail to induce a step movement into the cage. After a 2-minute rest, the procedure was repeated at a jump distance of 5 cm to allow the mouse to recognize that the inside of the cage was more comfortable than the jump platform. After another 2-minute rest, the jump distance was extended to 10 cm and the same hindquarter stimulus was applied to induce a jump. A jump was scored as success if both forelimbs reached the outer edge of the cage, and as a failure otherwise. When all four limbs reached the cage, the jump distance was recorded as the measured distance from the leading edge of the platform to the landing point of the forelimbs, rounded down to the nearest multiple of 5 cm. In the case of failure, the mouse was immediately returned to the jump platform without a rest period; in the case of success, the mouse was allowed to rest. After a 2-minute rest, the trial was repeated at a jump distance of 10 cm to allow the mouse to learn the jumping movement. The jump distance was then increased in 5-cm increments until 5 consecutive failures occurred, and the last successful jump distance was defined as the maximum jump distance.

### EMG ELECTRODE IMPLANTATION SURGERY

Only mice that jumped reliably in the jump test on Day 1 underwent surgery. This avoided operating on animals unlikely to yield usable recordings. EMG electrode implantation surgery was performed on 17 WT and 15 GOF mice that met this criterion (Fig. 1C). General anesthesia was induced by intraperitoneal injection of medetomidine, midazolam, and butorphanol (0.75, 4.0, and 5.0 mg/kg, respectively), and adequate depth of anesthesia was confirmed by the loss of the corneal reflex. An approximately 1.5 cm longitudinal skin incision was made from the forehead to the top of the head. The periosteum was dissected using forceps and a pointed scalpel and then air-dried. Next, the iliac crest was palpated, and an approximately 1.5 cm longitudinal skin incision was made along the dorsal midline from that level toward the head; the subcutaneous tissue was bluntly dissected with curved forceps to connect this wound to the head wound via a subcutaneous tunnel. An approximately 2-cm longitudinal skin incision was made from the lateral aspect of the left knee joint to the distal lateral lower leg, and the subcutaneous tissue was bluntly dissected to connect this wound to the dorsal wound via a subcutaneous tunnel.

To record muscle activity, pairs of Teflon-coated annealed stainless-steel wires (outer diameter, 0.14 mm; cat. no. 793200, A-M Systems, Sequim, WA, USA) were passed through a subcutaneous tunnel from the head to the lower leg. The superficial fascia of the lower leg was incised to expose the lateral aspects of the gastrocnemius and tibialis anterior muscles. Two electrodes were sutured into each of the gastrocnemius and tibialis anterior muscles at 5 mm intervals. A single electrode wire was sutured subcutaneously in the dorsal wound as a ground electrode. The electrode connector was fixed to the periosteum-exposed area of the skull using dental adhesive (Super-Bond, Sun Medical Co., Ltd., Shiga, Japan) and light-curing resin (Estelite Universal Flow Medium, Tokuyama Dental Corporation, Tokyo, Japan), completing the surgery. After completion of surgery, an antagonist (atipamezole, 1.5 mg/kg) was administered, and the mice were returned to cages containing paper-based bedding with high heat-retaining properties to keep them warm.

On Day 3, under inhalation anesthesia (sevoflurane 3%, flow rate 0.3 L/min), the wound was checked for dehiscence, and the electrode wires were checked for exposure, and it was confirmed that the recording cable could be attached to and detached from the head connector. After attaching and detaching the cable, the fixation of the head connector was reinforced with sutures.

### EMG RECORDING

Before EMG recording, mice performed practice jumps without the cable attached. After confirming that jumping could be induced, the recording cable was attached to the head connector while the mouse was briefly sedated with inhalation anesthesia (sevoflurane 3%, flow rate 0.3 L/min). Before starting the recording, sufficient recovery of consciousness was confirmed, and EMG was recorded once no residual sedative effect remained. Measurements were performed starting at a jump distance of 5 cm and increasing in 5-cm increments, up to the maximum jump distance. EMG was measured up to the maximum jump distance based on visual confirmation of 8 successful jumps as a guide, and in some mice, EMG was also measured at specific distances such as 15 cm, 25 cm, and 35 cm. EMG signals were amplified (x500) and band-pass filtered at 15–10 kHz with an amplifier (MEG-6108MMG, MIYUKI GIKEN Co., Ltd., Tokyo, Japan) and digitized at 10 kHz using PowerLab (2602-0301, ADInstruments, Dunedin, New Zealand) with LabChart software (v8.1.16) (Fig. 1D).

### COMPOUND MUSCLE ACTION POTENTIAL RECORDING

To compare EMG amplitudes across animals, CMAP was measured immediately after the jumping experiment (Fig. 1E). To avoid displacement of the electrode position, mice were sedated with inhalation anesthesia and then anesthetized by intraperitoneal injection of the same anesthetic mixture used for surgery. As described by Arnold et al. (2015), two needle electrodes (300 μm diameter, an inter-electrode distance of 5 mm, #300UM, BioResearchCenter, Tokyo, Japan) were inserted percutaneously over the left sciatic nerve at the buttock region. Electrical stimulation was applied using an electrical stimulator (USE-210-S5, Unique Medical Co., Ltd., Tokyo, Japan) at intensities of 0.5, 1.0, 1.2, 1.5, 2.0, 3.0, and 5.0 times the movement threshold intensity at a fixed pulse width (0.1 ms). At each intensity, 20 stimuli were delivered at 1 Hz. Stimulus-evoked responses were recorded from gastrocnemius and tibialis anterior muscles with the same implanted electrodes, using the same recording system and settings described above, over an epoch extending from 2 ms before to 10 ms after each stimulus.

### EMG SIGNAL PROCESSING

The EMG signals were band-pass filtered (30–1500 Hz, second-order, zero-lag Butterworth filter), full-wave rectified, and low-pass filtered at 50 Hz (third-order, zero-lag Butterworth filter) to obtain the EMG envelope (Fig. 1D). To compensate for inter-animal differences in electrode placement and tissue impedance, EMG amplitudes were normalized to the maximal CMAP amplitude (Zellers et al., 2019). Peak-to-peak CMAP amplitudes were calculated and averaged across the 20 stimuli delivered at each intensity. The resulting recruitment curve was fitted with a four-parameter logistic function (Klimstra and Zehr, 2008; Fig. 1E), and the upper asymptote of the fitted curve was defined as the maximal CMAP amplitude used for EMG normalization. Fit quality was assessed by the coefficient of determination (R²), which exceeded 0.9 for all animals and both muscles. Normalization was performed separately for each muscle in each animal. Jump epochs were extracted from 150 ms before to 50 ms after takeoff and aligned to takeoff (Fig. 1D). For each animal, the peak amplitude and the full width at half maximum (FWHM) of the normalized EMG envelope were quantified for each jump and averaged across the 5–6 jumps at each jump distance.

### VIDEO RECORDING

Jumps performed during EMG recording were captured with three cameras (GoPro HERO10 Black; GoPro, San Mateo, CA, USA) at 240 frames/s and 1920 × 1080 pixel resolution. One camera was positioned overhead (overhead camera), one at foot level lateral to the platform (near-lateral camera), and one laterally at foot level but set back from the landing point to capture the entire jumping motion (set-back lateral camera). Video and EMG recordings were synchronized by TTL pulses generated by the PowerLab, which drove an LED visible to all three cameras. Jump success was verified by slow-motion playback, and the landing position was determined from the overhead camera to confirm the actual jump distance. Takeoff was defined as the first frame in which both hindlimbs had lost contact with the platform, identified from the near-lateral camera by frame-by-frame inspection (Fig. 1D).

#### Markerless tracking and ankle angle measurement

To quantify hindlimb kinematics during jumping, we measured the two-dimensional positions of the hindlimb joints using DeepLabCut (version 3.0.0rc9, PyTorch engine) (Mathis et al., 2018; Nath et al., 2018). Tracking was performed on video from the set-back lateral camera, which captured the entire jumping motion. For each jump, a window from 300 ms before to 50 ms after takeoff (84 frames) was extracted for analysis.

First, we annotated the two-dimensional positions of the knee, the ankle, and the foot of the tracked hindlimb, together with two static markers on the beam (beam1 and beam2) that served as spatial references and were not used in the angle computation (Fig. 1D).

Next, we trained a deep neural network based on transfer learning of the pre-trained network (ResNet50). The training set was built in two stages. We first labeled a total of 240 images, comprising 20 evenly spaced frames from each of 12 jumps (6 jumps each from one WT and one GOF mouse), so that variation across repeated jumps within an animal was represented before variation between animals was introduced. We then applied the trained network to one jump from each of the remaining 30 mice, corrected the predicted labels by hand, added them to the training set and retrained the network. This predict–correct–retrain cycle was repeated until all animals were represented, giving a final training set of 1,053 labeled images from 42 jumps in 32 mice. 95% of the labeled frames were used for training and the remaining 5% were held out for testing, and the network was trained for 200 epochs; the best-performing snapshot on the test set, at epoch 110, was used for all subsequent predictions.

The network output was used only to pre-position the keypoints. The trained network was applied to every analyzed recording, and every keypoint in every analyzed frame was then inspected and repositioned by hand in the DeepLabCut labelling GUI. All angles reported here were therefore computed from manually placed labels.

We then used the manually placed coordinates to calculate the ankle angle in each frame, defined as the angle subtended at the ankle between the foot and the knee (dorsiflexion decreases this angle). The angle time series of each jump was low-pass filtered at 20 Hz (third-order, zero-lag Butterworth filter). The angles were analyzed over frames 36–75, that is, 40 consecutive frames from −150 ms to +12.5 ms relative to takeoff (Fig. 1D). The window matched the onset of the EMG epoch and ended when the knee (the first landmark lost from view) was no longer visible in all trials. For each jump, the minimum ankle angle within this window was taken as the maximum dorsiflexion, and these values were averaged across the same 5–6 jumps used for the EMG analysis.

To examine when the genotype difference in muscle activity arose, the normalized EMG envelope of each jump was realigned to the time of maximum dorsiflexion of that jump, and mean envelope amplitude was measured over a pre epoch (−60 to −5 ms) and a post epoch (+5 to +30 ms); the 5 ms on either side was excluded because the time of maximum dorsiflexion could be resolved only to within one video frame (4.2 ms). Epoch values were obtained for each jump and averaged across the 5–6 jumps of each animal.

### TISSUE SAMPLING

On Day 8, the gastrocnemius muscle and Achilles tendon of the right hindlimb, which had not undergone electrode implantation, were collected and weighed. Tissues were collected only from the animals analyzed at the maximum jump distance. One WT mouse died from an undetermined cause between the EMG recording and tissue sampling, and no tissue was obtained from it; its EMG data were retained in the analysis. A predetermined subset of animals was processed for RNA extraction, and total RNA was extracted from each tissue using TRI Reagent (TR118, Molecular Research Center, Cincinnati, OH, USA). Body and gastrocnemius weights were not recorded for two WT mice, which were nonetheless included in the gene-expression analysis. One gastrocnemius sample from each genotype was excluded from the gene-expression analysis owing to insufficient RNA yield.

#### Real-time quantitative PCR

The extracted total RNA was reverse-transcribed into cDNA using the PrimeScript RT Reagent Kit (RR037B, Takara Bio, Tokyo, Japan). Real-time qPCR was performed using QuantStudio1 (A40427, Thermo Fisher Scientific, Waltham, MA, USA).

### STATISTICAL ANALYSES

Data are presented as mean ± SD unless otherwise stated. Comparisons between WT and GOF mice were performed using unpaired Welch’s *t* tests. To assess the effect of jumping distance in the representative animal shown in Fig. 2, one-way analysis of variance (ANOVA) followed by Tukey’s multiple comparison test was performed. To examine when the genotype difference in muscle activity arose (Fig. 5), three-way mixed ANOVA was performed with jump condition (maximal, submaximal) and genotype (WT, GOF) as between-subject factors and epoch (pre, post) as a within-subject factor, followed by Šídák-corrected multiple comparisons between pairs differing in a single factor. EMG signals and kinematic data were processed using MATLAB R2021b (MathWorks, Natick, MA, USA), and statistical tests were performed using GraphPad Prism 11 (GraphPad Software, San Diego, CA, USA).

**Figure 2.**
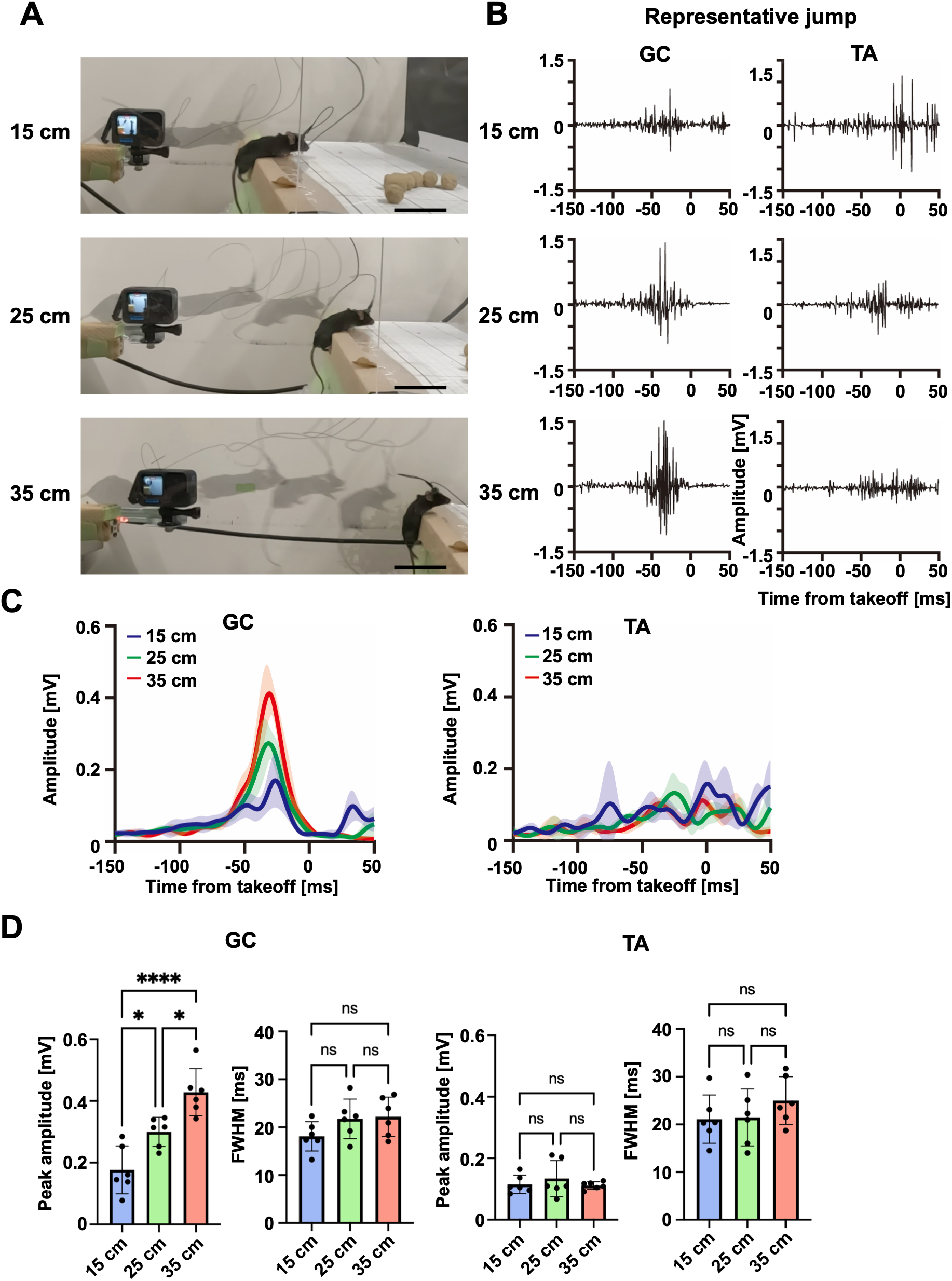
Different muscle activities at different jump distances. (A) Representative jumps from one WT mouse, from which all data in this figure were obtained. Top, 15 - cm jump; middle, 25-cm jump; bottom, 35-cm jump. Scale bar, 5 cm. (B) Representative raw electromyographic (EMG) traces from the same jumps shown in A, for the two hindlimb muscles: gastrocnemius (GC) and tibialis anterior (TA). (C) Mean EMG envelope at different jump distances (n = 6 jumps per distance). Shading indicates standard deviation (SD). (D) Peak amplitude and full width at half maximum (FWHM) of the EMG envelope for each muscle. Bars and error bars indicate mean ± SD; dots indicate individual jumps. \**p* < 0.05; \*\*\*\**p* < 0.0001; ns, not significant (Tukey test).

### DATA AND CODE ACCESSIBILITY

The MATLAB code used for EMG and kinematic analysis is available from the corresponding author upon reasonable request. The data reported in this study are available from the corresponding author upon reasonable request.

## Results

### Relationship between jump distance and hindlimb EMG

We first examined how hindlimb muscle activity varies with jump distance (Fig. 2A). In a representative WT mouse performing 6 jumps at each of 15, 25, and 35 cm, gastrocnemius (GC) activity increased progressively with jump distance, whereas tibialis anterior (TA) activity showed no distance-dependent change (Fig. 2B, C). Gastrocnemius peak amplitude increased significantly with distance (Fig. 2D; 0.18 ± 0.08, 0.30 ± 0.05, and 0.43 ± 0.08 mV at 15, 25, and 35 cm; one-way ANOVA, *F*(2,15) = 20.31, *p* < 0.0001; Tukey’s post hoc test, 15 vs 35 cm, *p* < 0.0001; 25 vs 35 cm, *p* = 0.014; 15 vs 25 cm, *p* = 0.018). In contrast, gastrocnemius FWHM did not differ significantly (Fig. 2D; 18.1 ± 3.1, 21.7 ± 4.1, and 22.2 ± 4.1 ms; one-way ANOVA, *F*(2,15) = 2.11, *p* = 0.16). In the tibialis anterior, neither the peak amplitude (Fig. 2D; 0.11 ± 0.03, 0.13 ± 0.06, and 0.11 ± 0.01 mV; one-way ANOVA, *F*(2,15) = 0.59, *p* = 0.57) nor the FWHM (Fig. 2D; 21.1 ± 5.1, 21.5 ± 6.0, and 25.0 ± 5.0 ms; one-way ANOVA, *F*(2,15) = 0.96, *p* = 0.41) differed significantly with distance. We thus confirmed that gastrocnemius activity scaled in amplitude with jump distance, without a change in burst duration or in tibialis anterior activity.

### Between-group comparison of WT and tendon-specific *Piezo1* GOF mice

We next asked what distinguishes maximal jumping between the two genotypes. Because only gastrocnemius activity scaled with jump distance, we compared its EMG at maximum jump distance (Fig. 3A). Jump distance and EMG were obtained from 12 WT and 10 GOF mice. As in the previous report (Nakamichi et al., 2022), the maximum jump distance was significantly longer in GOF mice than in WT mice (Fig. 3B; 46 ± 4 cm in WT vs 50 ± 5 cm in GOF; Welch’s *t* test, *t*(18.23) = 2.17, *p* = 0.043). Notably, the gastrocnemius burst was larger in GOF than in WT mice (Fig. 3C, D). Peak amplitude normalized to CMAP was significantly higher in GOF mice (Fig. 3E; 2.4 ± 0.5% in WT vs 3.3 ± 0.6% in GOF; Welch’s *t* test, *t*(16.24) = 3.62, *p* = 0.0023). Because amplitudes were normalized to the maximal CMAP of each animal (Fig. 1E), this difference is unlikely to reflect electrode placement or muscle size. This difference was maintained when the comparison was restricted to the distances at which both genotypes were represented (45 and 50 cm; 9 WT and 6 GOF mice), where distance itself did not differ significantly (Fig. S2A; 48 ± 3 cm in WT vs 47 ± 3 cm in GOF; Welch’s *t* test, *t*(11.03) = 0.81, *p* = 0.44) but peak amplitude remained higher in GOF mice (Fig. S2B; 2.5 ± 0.4% in WT vs 3.3 ± 0.7% in GOF; Welch’s *t* test, *t*(7.49) = 2.37, *p* = 0.047), suggesting that the amplitude difference was not explained by the longer jumps of GOF mice alone. The FWHM, in contrast to the peak amplitude, did not differ significantly (Fig. 3E; 21.1 ± 2.8 ms in WT vs 19.4 ± 1.8 ms in GOF; Welch’s *t* test, *t*(18.86) = 1.71, *p* = 0.10). Thus, during maximal jumping, the two genotypes differed in the amplitude, but not the duration, of gastrocnemius activity.

**Figure 3.**
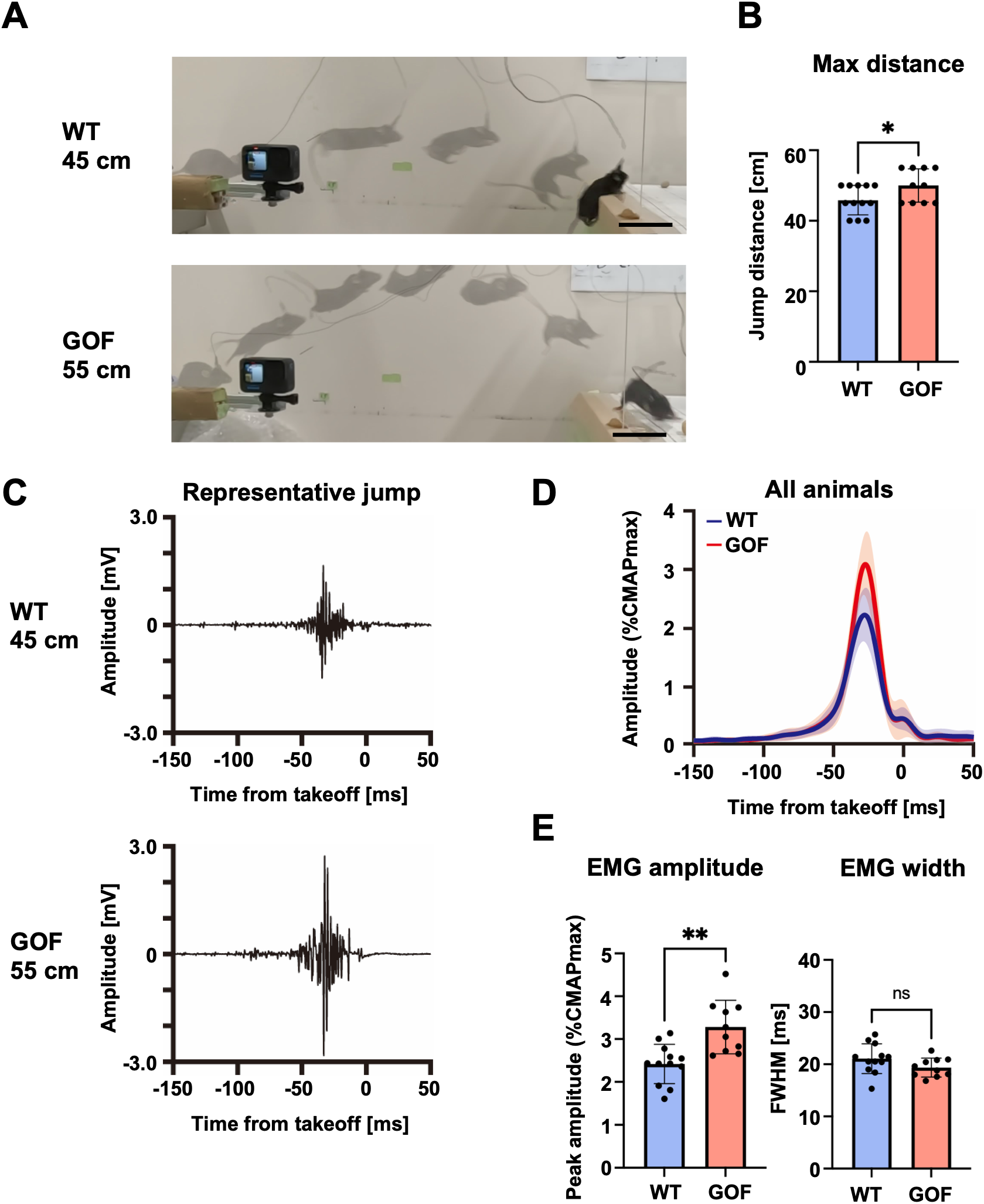
Different muscle activities between WT and GOF mice at maximum jump distances. (A) Representative jumps at the maximum jump distance. Top, WT (45 cm); bottom, GOF (55 cm). Scale bar, 5 cm. (B) Maximum jump distance. Bars and error bars indicate mean ± SD; dots indicate individual animals. \**p* < 0.05 (Welch’s *t* test). (C) Representative raw gastrocnemius EMG traces from the same jumps shown in A. (D) Mean gastrocnemius EMG envelope of WT and GOF mice (WT, n = 12; GOF, n = 10), normalized to the maximum CMAP. Shading indicates SD. (E) Peak amplitude and FWHM of the gastrocnemius EMG envelope. Bars and error bars indicate mean ± SD; dots indicate individual animals. \*\**p* < 0.01; ns, not significant (Welch’s *t* test).

We next asked whether this elevation of gastrocnemius activity is specific to the maximum jump distance or is instead a constitutive feature of GOF mice. We therefore compared the gastrocnemius EMG during 25-cm jumps, a distance that is submaximal for both genotypes. EMG was obtained from 5 WT and 5 GOF mice (Fig. 4A, B). CMAP-normalized peak amplitude did not differ significantly between genotypes (Fig. 4C; 2.1 ± 0.3% in WT vs 2.1 ± 0.5% in GOF; Welch’s *t* test, *t*(7.11) = 0.17, *p* = 0.87), nor did the FWHM (Fig. 4C; 21.9 ± 4.3 ms in WT vs 20.4 ± 2.0 ms in GOF; Welch’s *t* test, *t*(5.73) = 0.70, *p* = 0.51). Thus, the enhanced gastrocnemius output in GOF mice was specific to the maximum jump distance, the highest effort level each animal could produce. A general increase in neural drive to the gastrocnemius would have been expected at every distance; its absence at 25 cm indicates that the enhancement depended on load.

**Figure 4.**
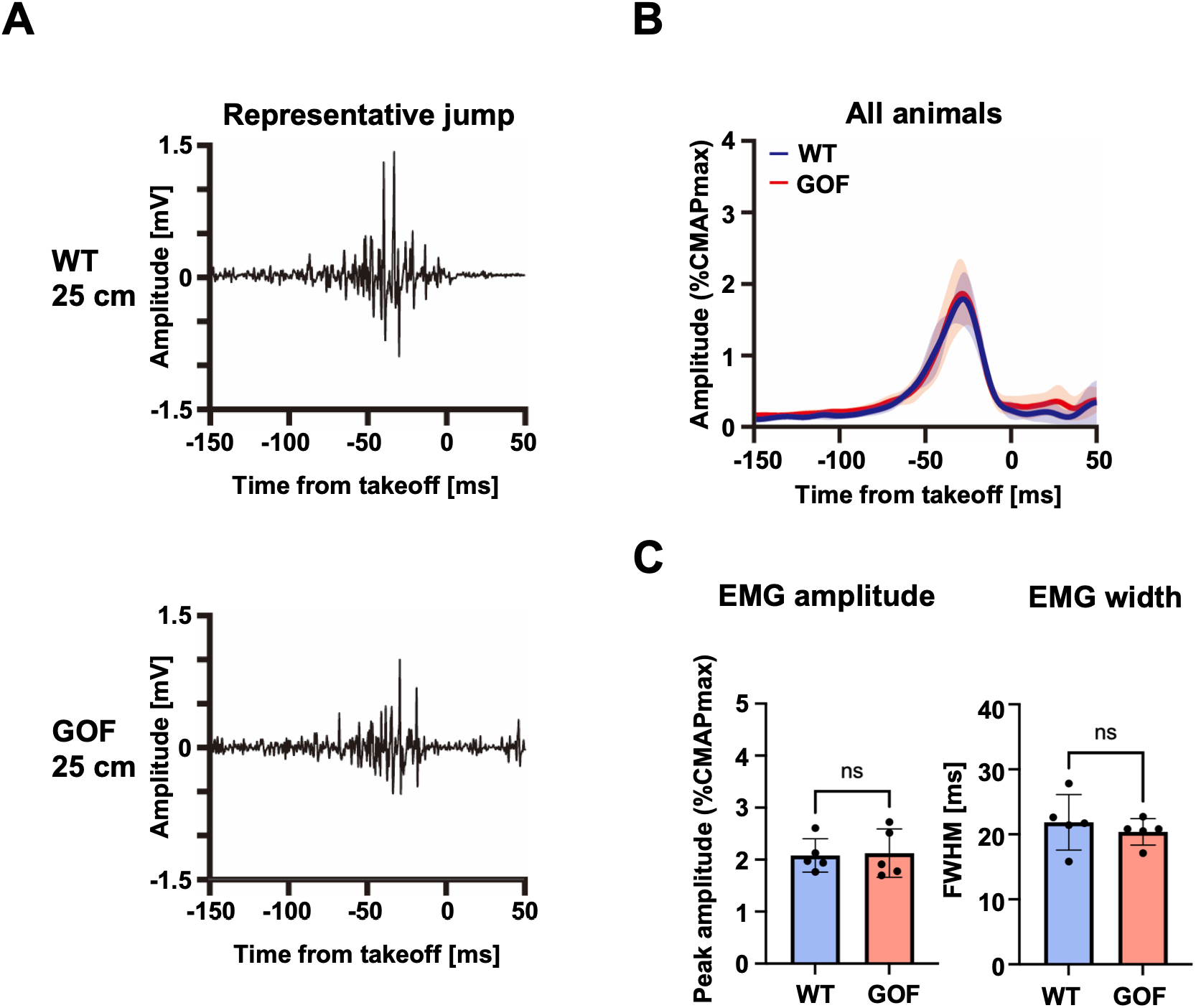
Comparable muscle activities between WT and GOF mice at submaximal jump distance. (A) Representative raw gastrocnemius EMG traces from a single 25-cm jump in one WT and one GOF mouse. (B) Mean gastrocnemius EMG envelope of WT and GOF mice (n = 5 per genotype), normalized to the maximum CMAP. Shading indicates SD. (C) Peak amplitude and FWHM of the gastrocnemius EMG envelope. Bars and error bars indicate mean ± SD; dots indicate individual animals. ns, not significant (Welch’s *t* test).

**Figure 5.**
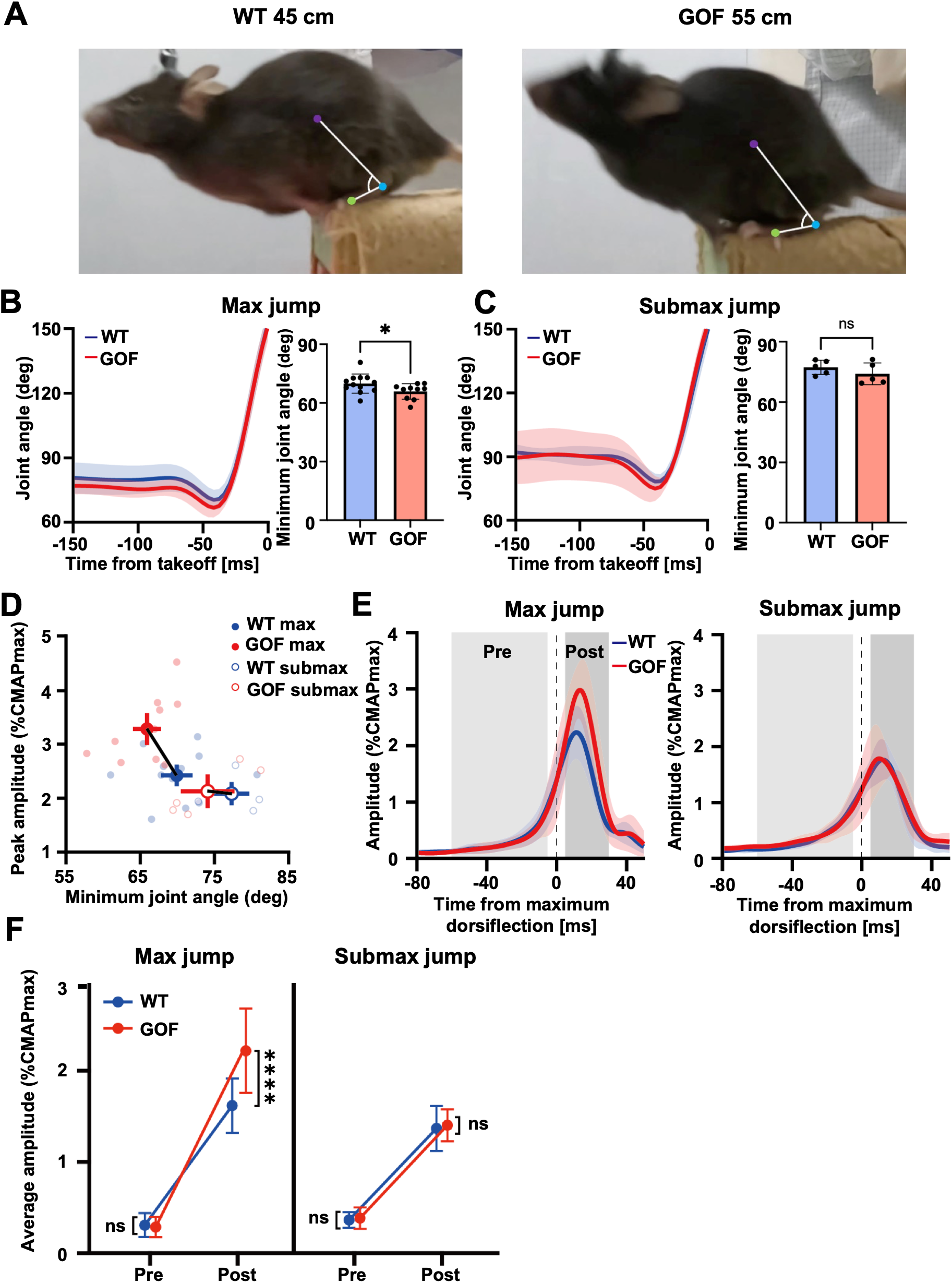
Enhanced EMG in GOF mice emerges after maximum dorsiflexion (A) Representative video frames at the minimum ankle joint angle during maximal jumping. Left, WT (45-cm jump; minimum angle 72°); right, GOF (55-cm jump; 65°). Purple dots indicate the knee, blue dots the ankle, and green dots the foot; the arc denotes the ankle joint angle. (B) Left, group-averaged time course of the ankle joint angle during maximal jumps (WT, n = 12; GOF, n = 10). Solid lines indicate the group mean; shading indicates SD. Right, minimum ankle joint angle during maximal jumping for each animal. Bars and error bars indicate mean ± SD; dots indicate individual animals. \**p* < 0.05 (Welch’s *t* test). (C) Left, group-averaged time course of the ankle joint angle during submaximal jumps (n = 5 per genotype). Solid lines indicate the group mean; shading indicates SD. Right, minimum ankle joint angle during submaximal jumping for each animal. Bars and error bars indicate mean ± SD; dots indicate individual animals. ns, not significant (Welch’s *t* test). (D) Peak normalized gastrocnemius EMG amplitude plotted against the minimum ankle joint angle for each animal at the maximal and submaximal jump distances (maximal jumps: WT, n = 12; GOF, n = 10; submaximal jumps: n = 5 per genotype). Each symbol represents one animal. Crosses indicate the group mean ± SEM in both dimensions. (E) Group-averaged time course of the normalized gastrocnemius EMG amplitude. Left, maximal jumps (WT, n = 12; GOF, n = 10); right, submaximal jumps (25 cm; n = 5 per genotype). Solid lines indicate the group mean; shading indicates SD. The dashed line indicates 0 ms, the time at which the ankle joint angle reached its minimum (maximum dorsiflexion). Light gray background shading marks the Pre epoch (−60 to −5 ms) and dark gray background shading marks the Post epoch (+5 to +30 ms), the two windows compared in F. (F) Normalized gastrocnemius EMG amplitude in the Pre and Post epochs defined in E. Left, maximal jumps; right, submaximal jumps. Symbols indicate the group mean; error bars indicate SD. \*\*\*\**p* < 0.0001; ns, not significant (Šídák-corrected comparisons following three-way mixed ANOVA).

### Relationship between ankle joint kinematics and gastrocnemius EMG

The preceding results showed that gastrocnemius output was enhanced in GOF mice only at maximum jump distance. To determine what distinguishes the two genotypes, we examined ankle joint kinematics, measuring the ankle angle during jumping (Fig. 5A). At maximum jump distance, the minimum ankle angle — that is, the deepest dorsiflexion reached before takeoff — was significantly smaller in GOF than in WT mice (Fig. 5B; 69.9 ± 4.9° in WT vs 65.9 ± 4.0° in GOF; Welch’s *t* test, *t*(20.0) = 2.13, *p* = 0.046). At the submaximal distance, the two genotypes did not differ significantly (Fig. 5C; 77.3 ± 3.5° in WT vs 74.1 ± 5.4° in GOF; Welch’s *t* test, *t*(6.91) = 1.12, *p* = 0.30). Thus, the minimum ankle angle only differentiated the two genotypes at maximum jump distance, which is the same condition in which the EMG difference was found. The influence of the altered tendon may therefore emerge only when the muscle–tendon unit is more heavily loaded.

To display the two measures together, we plotted the peak gastrocnemius amplitude of each animal against its minimum ankle angle (Fig. 5D). At maximum jump distance, the GOF group mean lay at a smaller minimum angle and a larger peak amplitude than the WT mean, whereas at the submaximal distance the two genotypes coincided. Thus, two independent manipulations — genotype and jump condition — shifted the group means in the same direction, toward deeper dorsiflexion and greater muscle activity.

This shift does not reveal which of the two measures changed first. We therefore realigned the gastrocnemius EMG envelope of each jump to the time of maximum dorsiflexion, i.e., the point of zero angular velocity at which lengthening of the muscle–tendon unit ends, and compared the EMG amplitude in the epochs before (pre) and after (post) maximum dorsiflexion (Fig. 5E). A three-way mixed ANOVA showed that the increase in amplitude from the pre to the post epoch was significantly larger in GOF than in WT mice (epoch × genotype, *F*(1, 28) = 5.86, *p* = 0.022), and that the extent of this genotype effect itself depended on jump condition (epoch × jump condition × genotype, *F*(1, 28) = 5.33, *p* = 0.029). Post hoc comparisons showed that the genotype difference was observed only in the post epoch at maximum jump distance (1.6% in WT vs 2.2% in GOF; mean difference, 0.60%; 95% CI, 0.28‒0.93%; Šídák-corrected multiple comparisons, *t*(56) = 5.50, *p* < 0.0001), with no difference in the pre epoch at maximum jump distance or in either epoch at the submaximal distance (Fig. 5F). Because no genotype difference was present before maximum dorsiflexion, this enhancement is unlikely to reflect a difference in feed-forward descending drive already present at that time, and is instead consistent with an origin in feedback arising during the jump itself.

### Tissue evaluation

Finally, we performed tissue analyses to address the possibility that the change in muscle activity had arisen from the muscle itself. Body weight and gastrocnemius wet weight did not differ between genotypes (Fig. 6A, body weight, 27.2 ± 1.5 g in WT vs 27.4 ± 1.2 g in GOF; Welch’s *t*-test, *t*(15.3) = 0.25, *p* = 0.81; gastrocnemius weight, 119.4 ± 9.0 mg in WT vs 121.4 ± 10.0 mg in GOF; Welch’s *t*-test, *t*(17.00) = 0.47, *p* = 0.65), nor did expression of myogenic markers in muscle (Fig. 6B): *MyoD* (1.00 ± 0.58 in WT vs 0.67 ± 0.45 in GOF; Welch’s *t* test, *t*(7.51) = 1.02, *p* = 0.34), the slow-twitch marker *Myh7* (1.00 ± 0.55 in WT vs 0.93 ± 0.95 in GOF; Welch’s *t* test, *t*(6.39) = 0.13, *p* = 0.90), or the fast-twitch marker *Myh1* (1.00 ± 0.11 in WT vs 1.52 ± 0.99 in GOF; Welch’s *t* test, *t*(4.10) = 1.17, *p* = 0.31). By contrast, expression of tendon markers was higher in GOF mice (Fig. 6C): *Mkx* (1.00 ± 0.50 in WT vs 4.27 ± 1.67 in GOF; Welch’s *t* test, *t*(5.90) = 4.59, *p* = 0.0039), *Scx* (1.00 ± 0.56 in WT vs 4.18 ± 2.77 in GOF; Welch’s *t* test, *t*(5.41) = 2.75, *p* = 0.037), *Tnmd* (1.00 ± 0.35 in WT vs 2.08 ± 0.95 in GOF; Welch’s *t* test, *t*(6.35) = 2.59, *p* = 0.039), confirming that the manipulation acted on tendon. GTO morphology is also unchanged in this line (Nakamichi et al., 2022). Thus, the increased muscle activity during jumping in GOF mice is unlikely to be of muscular or receptor origin, pointing instead to a neural origin. Taken together, our results demonstrate that a genetic manipulation confined to tendon can shape neuromuscular output during ballistic movement.

**Figure 6.**
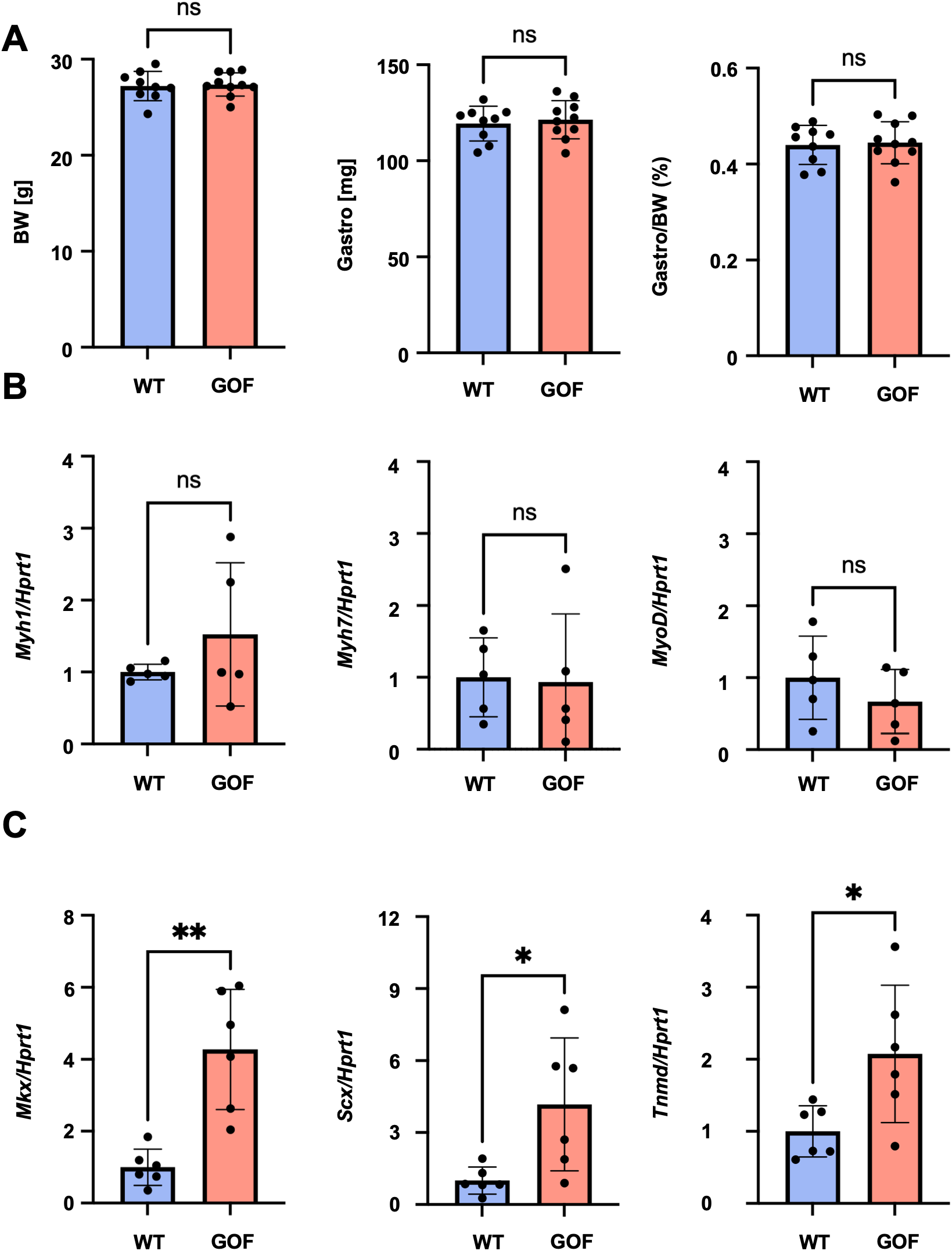
No difference in muscle mass or myogenic gene expression between WT and GOF mice. (A) Body weight (BW), gastrocnemius muscle wet weight, and gastrocnemius weight as a percentage of body weight in WT and *Piezo1* GOF mice (WT, n = 9; GOF, n = 10). Bars and error bars indicate mean ± SD; dots indicate individual animals. (B) Relative mRNA expression of *Myh1*, *Myh7*, and *MyoD* in gastrocnemius muscle (n = 5 per genotype). Expression was normalized to *Hprt1*. Bars and error bars indicate mean ± SD; dots indicate individual animals. (C) Relative mRNA expression of *Mkx*, *Scx*, and *Tnmd* in tendon (n = 6 per genotype). Expression was normalized to *Hprt1*. Bars and error bars indicate mean ± SD; dots indicate individual animals. \**p* < 0.05; \*\**p* < 0.01; ns, not significant (Welch’s *t* test).

## Discussion

Tendon, a metabolically inert and structurally stable tissue, has received far less attention than muscle or neural circuits in the field of motor control studies. We asked whether a change in mechanical properties confined to tendon alters the neural drive to muscle during movement. We compared hindlimb EMG and ankle kinematics between WT and tendon-specific *Piezo1* GOF mice during jumping and obtained two main findings. First, gastrocnemius activity was larger in GOF than in WT mice at maximal jump but not at a submaximal distance, with no change in burst duration, muscle mass, or myogenic gene expression (Fig. 3E, 4C, 6A, B). Second, this genotype difference was absent before the ankle reached maximum dorsiflexion and emerged only afterwards (Fig. 5E, F). These findings indicate that a structural intervention confined to tendon altered neuromuscular output in a load-dependent manner.

Because EMG records the discharge of motor neurons, the genotype difference in EMG amplitude indicates a difference in neural drive. As shown in Results, this difference is not accounted for by muscle mass or fiber type, by GTO morphology, by electrode placement, or by the longer jumps of GOF mice (Fig. 6A, B; Fig. 1E; Fig. S2). A genetic manipulation confined to tendon therefore changed the neural drive to muscle: tendon mechanical properties act on motor output not only directly, through the series arrangement of tendon with muscle, but also indirectly, through the central nervous system. A purely mechanical account is insufficient: the gain in kinetic energy during jumping far exceeds the gain in elastic energy stored in the tendon (Nakamichi et al., 2022), so energy storage alone cannot explain how far GOF mice jump. The enhanced neural drive shown here supplies the missing component.

Which neural pathway could mediate this effect? The inputs that set the amount of muscle activity fall into two categories. One is descending motor commands from the brain, i.e., feedforward control (Evarts, 1968); the other is proprioceptive input from the moving limb, i.e., feedback control (Marsden et al., 1972; Wolpert and Bastian, 2021). Proprioceptive input itself arises from two receptors: muscle spindles, which lie within the muscle belly and give rise to Ia and group II afferents, and the GTO, which lies at the muscle–tendon junction and gives rise to Ib afferents. Here, we consider each of these possibilities.

A difference in feedforward drive is unlikely to account for our results. If GOF mice had received a larger descending command, the genotype difference in gastrocnemius activity should have been present from the onset of the burst. Instead, the two genotypes did not differ in the epoch before maximum dorsiflexion, and the difference emerged only in the epoch after it (Fig. 5E, F). The additional drive in GOF mice therefore arose during the movement rather than being specified before it.

Among feedback pathways, the stretch reflex mediated by Ia afferents from muscle spindles is the first candidate, because studies of human jumping attribute the improvement with deeper squatting to spindle-mediated reflex support of the extensors (Burke et al., 1983; Anderson and Pandy, 1993). GOF mice did reach a deeper dorsiflexion (Fig. 5B), so a spindle origin deserves consideration. Three features of our results, however, argue against this account. First, the modified tissue is the tendon, whereas spindles lie within the muscle belly. Second, Ia afferents are sensitive to the velocity of muscle stretch, yet the genotype difference emerged only after the point of zero angular velocity (Fig. 5E, F). Third, greater series compliance reduces the stretch transmitted to the muscle fibers for a given change in muscle–tendon unit length, and artificially increased series compliance attenuates spindle firing (Abbott et al., 2024); a spindle-mediated account therefore predicts a change in the opposite direction. The Ia pathway is thus an unlikely source of the enhanced drive.

The Ib pathway from the GTO, in contrast, fits each of these features: the GTO lies in the modified tissue, and Ib afferents signal force, not velocity. Ib input is also the only candidate that predicts the load dependence of our results. In the walking cat, Ib afferents from ankle extensors provide positive force feedback to the extensor motor pool, and the gain of this feedback increases with both the force on the tendon and the length of the muscle–tendon unit (Donelan and Pearson, 2004; Donelan et al., 2009). A pathway whose gain grows with load would contribute little at a submaximal distance and most at the maximal jump, which is the pattern we observed (Fig. 3E, 4C). That Ib input excites, rather than inhibits, the homonymous motor neurons during movement is established for locomotion (Guertin et al., 1995; Donelan et al., 2009), and we recently showed that the same excitatory loop contributes to muscle activity during voluntary hand movement in primates (Kim et al., 2026). The excitatory Ib loop thus operates across modes of movement and across species, from locomotion and jumping in quadrupeds to skilled hand use in primates, and is the most plausible source of the enhanced drive in GOF mice.

If the Ib pathway is engaged, how does a change confined to tendon increase its output? PIEZO1 transduces mechanical stress into structural remodeling of tendon, and in tendon-specific *Piezo1* GOF mice the collagen is reorganized and the tendon is more compliant, whereas the GTO itself is morphologically unchanged (Saotome et al., 2018; Nakamichi et al., 2022). Because the GTO is arranged in series within the muscle–tendon unit, the strain it experiences at a given muscle force depends on the compliance of the surrounding collagen (Houk and Henneman, 1967; Jami, 1992), as shown for the spindle (Abbott et al., 2024). We assume that the GOF tendon differs only in this compliance. During the countermovement, when the extensors contract while the ankle dorsiflexes, the more compliant tendon elongates more under the same force, which is seen as the deeper dorsiflexion of GOF mice (Fig. 5B), and the GTO in series with it would be strained more, so that Ib discharge for a given force would be larger. Two features of maximal jumping amplify this effect: the force on the tendon is greatest, as GOF mice jumped farther with no change in burst duration (Fig. 3E), and the muscle–tendon unit is longest, both of which raise the gain of Ib feedback (Donelan and Pearson, 2004). This larger Ib input, acting through the excitatory loop, then adds drive to the gastrocnemius motor pool. This sequence also specifies when the genotype difference should appear not at burst onset, but only after the tendon has been loaded, i.e., after maximum dorsiflexion. This is what we found (Fig. 5E, F). A change in tendon alignment therefore acts not only as a mechanical element of the muscle–tendon unit but also as a stimulus to the nervous system.

The account above assumes that the GOF mutation acted on the tendon alone. This assumption is supported on two grounds: expression of the mutant allele is restricted to *Scx*-expressing cells of tendon and ligament (Fig. S1), and the mechanotransduction channel of proprioceptors is PIEZO2, not PIEZO1 (Woo et al., 2015), so the mutation is unlikely to have altered the sensory neurons themselves. However, the tendon in these mice has been altered since development, and we cannot exclude an adaptation of the nervous system to this altered tendon that contributed to the enhanced drive through a route other than the one proposed here. Tamoxifen-inducible *Scx*-CreERT2 *Piezo1* GOF mice show tendon hypertrophy and improved jumping 12 weeks after adult induction (Nakamichi et al., 2022), but neuromuscular activity was not assessed in that line. Combining adult induction with the recordings used here would separate an acute effect of tendon compliance on Ib feedback from a chronic reorganization of the circuit.

A second limitation is that our inference rests on EMG and kinematic measures alone. We did not record Ib afferent activity, and the contributions of descending, Ia, and Ib inputs cannot be separated from EMG, so the Ib origin proposed here remains the most parsimonious account rather than a demonstrated one.

Although technical barriers remain, future studies should aim to test it directly by (i) recording Ib afferent discharge during jumping in the two genotypes, (ii) manipulating Ib afferents selectively, using genetic access based on candidate Ib markers such as *Pcdh8* (Oliver et al., 2021) and *Chad* (Wu et al., 2021), and (iii) manipulating descending pathways during the same task.

In summary, this study shows that a change in mechanical properties confined to tendon alters the neural drive to muscle during ballistic movement, and the pattern of this change points to Ib positive feedback from the GTO as its source. The same relationship may operate in the opposite direction with age: tendons stiffen with age (Heinemeier et al., 2013; Stowe et al., 2024), so attenuated Ib feedback could be one factor in the age-related decline of motor function. These findings suggest that tendon is not a passive transmitter of force but a sensory organ whose mechanical properties set the gain of the feedback that shapes motor output.

## Author Contribution

K.N., A.K., R.K., K.S. and H.A. designed research; K.N., A.K., R.N., R.K., T.Matsushima, T.C., M.K. and T.Morita performed research; K.N. and A.K. analyzed data; K.N., A.K., R.N., T.Y., K.S. and H.A. edited the paper; K.N., A.K, K.S. and H.A. wrote the paper.

The authors declare no competing financial interests.

## Acknowledgements

This work was supported by Japan Society for the Promotion of Science (JSPS) KAKENHI (Grant Numbers JP26K21770 and JP26K23559 to H.A., JP23H05488, 24K21313 to K.S.), Japan Agency for Medical Research and Development (AMED) (Grant Numbers JP22gm0010009, JP22ama121045, JP24gm2010002, JP25jf0126010 and JP256f0137003 to H.A.), Japan Science and Technology Agency (JST) GTIE GAP Fund (Grant Number JPMJSF2313 to H.A.), SPRING (Grant Number JPMJSP2180 to K.N.), CREST (Grant Number JPMJCR26F2 to K.S.), and Moonshot R&D(Grant Number JPMJMS263P to KS), and the JAXA Space Strategic Fund (SSF) Program Japan (Grant Number JPJXSSF25MX17005 to H.A.), and National Institutes of Health (NIH) (Grant Number R01AR080127 to H.A.)

The authors thank Chie Makino, Kasumi Yumoto, Maiko Inotsume and all the staff of the Department of Systems Biomedicine at Institute of Science Tokyo for their support and advice. Claude Opus 5.5 (Anthropic) was used for English translation and language editing of the manuscript. The authors reviewed all output and take full responsibility for the content. The schematics in Fig. 1B were created with BioRender (biorender.com) under a publication license.

---

**Figure S1.**
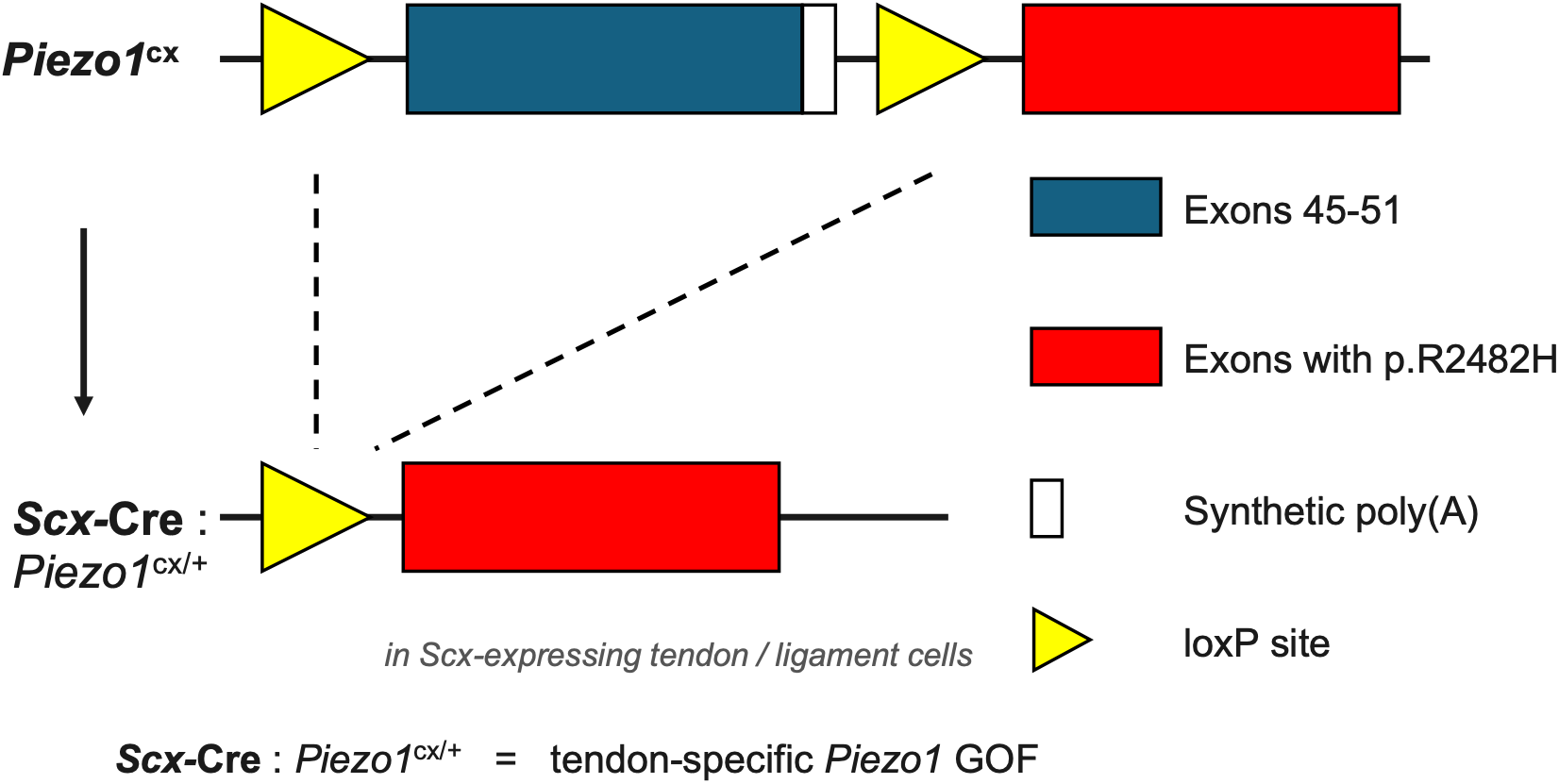
Targeting strategy for tendon-specific *Piezo1* gain of function. In the *Piezo1*^cx^ allele, loxP sites are inserted upstream and downstream of exons 45-51, and a second copy of exons 45-51 carrying the *Piezo1* GOF mutation p.R2482H is placed downstream of the 3′ loxP site, silenced by an intervening synthetic poly(A) cassette. Crossing *Piezo1*^cx^mice with a Cre line driven by *Scx*, a tendon-specific gene, excises the wild-type exons 45-51 and places the p.R2482H-containing exons, thereby generating tendon-specific *Piezo1* GOF mice.

**Figure S2.**
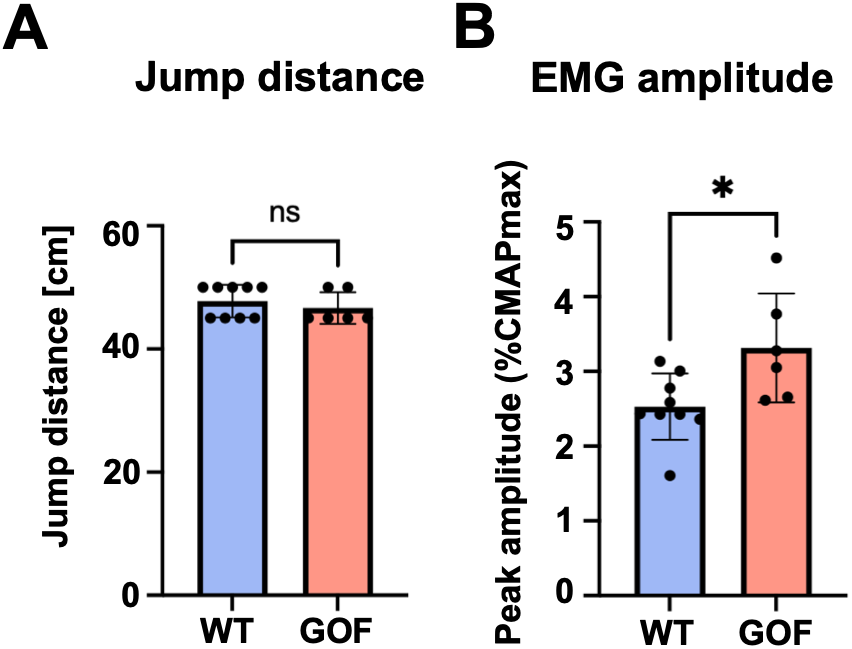
Different muscle activities between WT and GOF mice at comparable maximum jump distances. Analysis was restricted to animals whose maximum jump distance was 45 or 50 cm (WT, n = 9; GOF, n = 6) throughout. (A) Maximum jump distance. Bars and error bars indicate mean ± SD; dots indicate individual animals. ns, not significant (Welch’s *t* test). (B) Peak amplitude of the gastrocnemius EMG envelope, normalized to the maximum CMAP. Bars and error bars indicate mean ± SD; dots indicate individual animals. \**p* < 0.05 (Welch’s *t* test).

